# Manganese homeostasis modulates fungal virulence and stress tolerance in *Candida albicans*

**DOI:** 10.1101/2023.10.12.562042

**Authors:** Manon Henry, Inès Khemiri, Faiza Tebbji, Rasmi Abu-Helu, Antony T Vincent, Adnane Sellam

## Abstract

Due to the scarcity of transition metals within the human host, fungal pathogens have evolved sophisticated mechanisms to uptake and utilize these micronutrients at the infection interface. While considerable attention was turned to iron and copper acquisition mechanisms and their importance in fungal fitness, less was done regarding either the role of manganese (Mn) in infectious processes or the cellular mechanism by which fungal cells achieve their Mn- homeostasis. Here, we undertook transcriptional profiling in the pathogenic fungus *Candida albicans* experiencing both Mn starvation and excess to capture biological processes that are modulated by this metal. We uncovered that Mn scarcity influences diverse processes associated with fungal fitness including invasion of host cells and antifungal sensitivity. We show that Mn levels influence the abundance of iron and zinc emphasizing the complex crosstalk between metals. Deletion of *SMF12*, a member of Mn Nramp transporters confirmed its contribution to Mn uptake. *smf12* was unable to form hyphae and damage host cells and exhibited sensitivity to azoles. We found that the unfolded protein response (UPR), likely activated by decreased glycosylation under Mn limitation, was required to restore growth when cells were shifted from a Mn-starved to a Mn-repleted medium. RNA-seq profiling of cells exposed to Mn excess revealed that UPR was also activated. Furthermore, the UPR signaling axis Ire1-Hac1 was required to bypass Mn toxicity. Collectively, this study underscores the importance of Mn homeostasis in fungal virulence, and comprehensively provides a portrait of biological functions that are modulated by Mn in a fungal pathogen.

## Introduction

Transition metals such as iron (Fe), copper (Cu), zinc (Zn), and manganese (Mn) provide considerable functionality across biological systems as they are used as cofactors for catalytic enzymes and are thought to be required for the activity of one third of a cellular proteome (Waldron et al., 2009). Metal ions alter the physico-chemical properties of proteins, which promotes enzymatic catalysis, stabilization of protein structure and, electron and chemical group transfers (Murdoch and Skaar, 2022). At elevated concentrations, trace metals exhibit high toxicity which impose a tight regulation of their abundance at homeostatic levels. Trace metals are also deterministic for host-pathogen interaction as they are sequestered by the host to limit the proliferation of microbial pathogens, a process known as nutritional immunity (Hood and Skaar, 2012; Murdoch and Skaar, 2022). This process is achieved by the production of sequestering molecules such as calprotectin and siderocalin that chelate Mn, Zn and Fe (Sia et al., 2013; Hennigar and McClung, 2016; Zygiel and Nolan, 2018). Furthermore, ion metals are also sequestered in storage tissues and intracellular organelles which also limits the availability of those nutrient for uptake and use by invading pathogens (Hennigar and McClung, 2016; Murdoch and Skaar, 2022). Inversely, during infection, some immune cells tackle bacterial pathogens by releasing toxic levels of Cu or Zn which promotes microbial killing and attenuation of infectivity (Murdoch and Skaar, 2022).

As transition metal availability is very limited inside the human host, fungal pathogens have evolved sophisticated mechanisms to uptake and utilize these micronutrients at the infection interface. For instance, Fe acquisition by heme utilization and siderophore-mediated uptake are recognized as key virulence factors in many important fungal human pathogens including *Candida albicans*, *Aspergillus fumigatus* and *Cryptococcus neoformans* (Haas, 2012; Bairwa et al., 2017; Fourie et al., 2018; Roy et al., 2022). Similarly, Cu uptake is tightly controlled by the transcription factor Mac1 that is essential for fungal fitness *in vivo* (Cai et al., 2017; Culbertson et al., 2020; Khemiri et al., 2020). Zinc internalization by Zrts transporters or by the Zn scavenger Pra1 in *C. albicans* was also shown to be critical for the infectivity of this yeast and the expression of virulence traits (Citiulo et al., 2012; Gerwien et al., 2018; Soares et al., 2020). While considerable attention was turned to Fe, Cu and Zn acquisition mechanisms and their importance in fungal fitness, the role of Mn in infectious processes or the cellular mechanism by which fungal cells achieve their Mn-homeostasis is not well understood.

Mn is an essential trace element throughout the tree of life and serves as a cofactor for many enzymes such as polymerases, glycosyltransferases and superoxide dismutases (Martinez-Finley et al., 2013). Mammalian hosts restrict Mn availability by the Mn-chelating protein, calprotectin which contributes to antibacterial and antifungal defenses (Corbin et al., 2008; Brophy and Nolan, 2015; Besold et al., 2018). So far, the counteracting mechanisms to Mn sequestration by the host allowing Mn uptake and utilization by fungal pathogens remain unknown. In the budding yeast *Saccharomyces cerevisiae*, Mn uptake is achieved by Smf1 and Smf2 transporters of the NRAMP (natural resistance-associated macrophage protein) family in addition to the phosphate permease Pho84 which is also a low-affinity Mn transporter (Supek et al., 1996; Jensen et al., 2003; Culotta et al., 2005; Reddi et al., 2009; Cyert and Philpott, 2013). Unlike Fe, Zn and Cu transport that are subject to transcriptional control by metal-sensing transcription factors, Mn transport in *S. cerevisiae* is regulated at the post-translational level by ubiquitination and through changes in sub-cellular localization of Smf transporters (Culotta et al., 2005; Reddi et al., 2009; Cyert and Philpott, 2013). Under Mn sufficiency, Smf2 and Smf1 transit into the Golgi and are ubiquitinated by the E3 ubiquitin ligase Rsp5 which directs them to the vacuole for subsequent degradation, a process facilitated by adaptor proteins including Bsd2, Tre1 and Tre2 (Reddi et al., 2009). When Mn is depleted, Smf1 stability increases through conformation changes, which make it not recognizable by Rsp5/Bsd2, and subsequently localize to the plasma membrane for Mn uptake (Culotta et al., 2005). While the mechanisms preserving Mn homeostasis are well known in the saprophytic yeast *S. cerevisiae*, they remain largely unexplored in the rest of fungal species including human fungal pathogens. In *C. albicans*, deletion of *PHO84* had no impact on Mn uptake suggesting a diverging role for this transporter as compared to the budding yeast (Liu et al., 2018). Mn was also shown to be a critical cofactor for the *C. albicans* Mn-superoxide dismutases Sod2 and Sod3 and their contribution to oxidative stress defense (Rhie et al., 1999; Hwang et al., 2003; Wildeman et al., 2023).

In the current study, we undertook a transcriptional profiling of *C. albicans* cells experiencing both Mn starvation and excess to comprehensively capture biological processes that are modulated by Mn abundance. We uncovered that Mn scarcity influences diverse biological processes associated with fungal virulence including morphogenetic switch, metabolic flexibility, invasion of different host cells and antifungal and UPR stress responses. We also deleted one of the four Nramp transporters in *C. albicans*, named *SMF12* (Orf19.2270), and confirmed its contribution to Mn uptake and fungal fitness *in vivo*. While preparing this manuscript for submission, Wildeman *et al*. (Wildeman et al., 2023) published a similar work confirming the contribution of Smf12 to Mn homeostasis and its requirement for Mn-superoxide dismutase activity and oxidative stress resistance. We also analyzed the transcriptome of *C. albicans* cells exposed to Mn excess and uncovered that the UPR signaling was activated. Accordingly, we found that Ire1, the master regulator of UPR signaling in fungi, was essential to bypass Mn toxicity. Our RNA-seq analysis of the Mn-modulated transcriptome in *C. albicans* provides a global framework for future studies to examine the link between Mn metabolism and essential functions that modulate fungal virulence and fitness inside the host.

## Results and discussion

### Modulation of manganese content perturbs fungal growth

Due to the absence of a specific Mn chelator, we used a commercial Mn-free synthetic complete growth medium (SC-Mn) to assess the impact of Mn limitation on the growth of *C. albicans*. Quantification of Mn levels using ICP-MS confirmed the absence of Mn in the SC-Mn medium while showing the expected Mn amount in the conventional SC medium (SC) (**Figure 1A**). In the absence of Mn (SC-Mn), *C. albicans* exhibited a moderate growth reduction of roughly 20% as compared to the Mn replete condition (SC) (**Figure 1B**). Supplementation of SC-Mn medium with 1mM MnCl_2_ (SC+Mn) restored the growth of *C. albicans* to a level comparable to that observed in SC medium. To further modulate the intracellular levels of Mn, we created a mutant of one of the four predicted Mn transporters, Smf12 (orf19.2270) (**Figure 1C**). The ICP-MS analysis confirmed the low level of internalized Mn in *smf12* mutant under either Mn depletion (SC-Mn) or repletion conditions (SC+Mn or SC) suggesting that Smf12 is a major transporter of Mn in *C. albicans* (**Figure 1D**). Accordingly, *smf12* exhibited a significant growth defect in either Mn- repleted or depleted media as compared to the WT strain (**Figure 1E**). Taken together, these data suggest that modulation of intracellular Mn in *C. albicans* impacts the growth of this opportunistic yeast.

**Figure 1.**
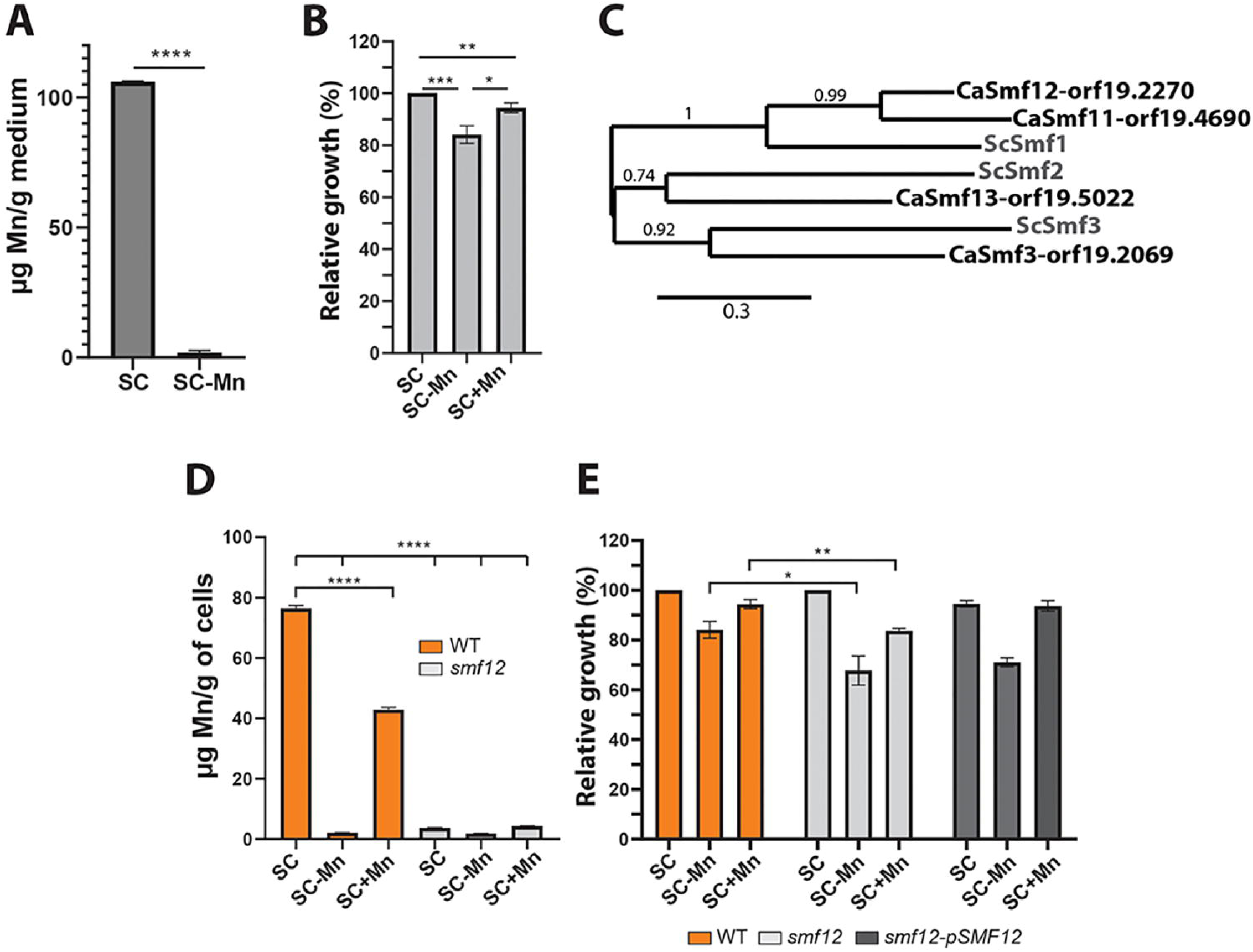
Modulation of manganese content perturbs fungal growth. (**A**) Mn quantification by ICP-MS in conventional synthetic complete (SC) and Mn-free SC media (SC-Mn). (**B**) Impact of Mn depletion on *C. albicans* growth. WT strain (SC5314) was grown at 30°C for 13 hrs in SC-Mn and SC+Mn media. Growth was normalized to the SC condition. (**C**) Phylogram of Smf transporters in *C. albicans* (CaSmf11-orf19.4690, CaSmf12-orf19.2270, CaSmf13-orf19.5022 and CaSmf3-orf19.2069) and *S. cerevisiae* (ScSmf1, ScSmf2 and ScSmf3). The multiple sequence alignment for the four Smf orthologues and the phylogenetic tree were generated using Clustal Omega. The branch length is proportional to the number of substitutions per site. (**D**) Mn uptake in the *smf12* mutant. Mn levels were determined by ICP-MS in both exponentially grown WT and *smf12* strains in SC, SC-Mn, and SC+Mn media at 30°C. (**E**) Impact of Mn depletion on *smf12* mutant. WT strain (SN250), *smf12* and the complemented *smf12* (s*mf12*- p*SMF12*) strains were grown at 30°C for 13 hrs in SC-Mn or SC+Mn. Growth was normalized to the SC condition. Statistics are multiple t-test with **** p-value < 0.00001, *** p-value < 0.0001, ** p-value < 0.001, * p-value < 0.05.

### Genome wide transcriptional response of *C. albicans* to Mn starvation

Although Mn is an important metal for different microorganisms the overall impact of its limitation on fungal biology is not well characterized so far. To define the cellular processes that are modulated by Mn in the pathogenic yeast *C. albicans*, we monitored the global gene expression dynamic under Mn insufficiency using RNA-seq. The transcriptome of cells growing in SC-Mn was compared to that of cells thriving in SC+Mn at 5 and 90 min to capture early cellular reprograming and steady-state adaptive mechanisms of Mn starvation, respectively. Gene ontology analysis revealed that Mn limitation at 5 and 90 min induces significant metabolic reprogramming as reflected by the upregulation of transcripts related to carbohydrate transport, galactolysis, carnitine biosynthesis and, fatty acid and nitrogen utilization (**Figure 2A-B and Supplementary Table S1**). The transcriptional activation of metabolic genes might reflect a compensatory response by *C. albicans* cells to sustain their metabolic needs as Mn is required for many metabolic metalloenzymes such as transferases, dehydrogenases and oxidases (Weatherburn, 2000). The 90 min time point was notable for the upregulation of genes involved in iron transport (*FRE7*, *FRE30*, *CFL2,4,5*, *FTH1*, *RBT5*, *SIT1*) and the downregulation of transcripts of mannosyltransferases (*MNN41*, *MNN42*, *BMT7*) and cell wall integrity proteins (*ECM331*, *PGA23*) (**Figure 2C-D**). Genes modulated by Mn include different virulence factors required for the yeast-to-hyphae transition (*HWP1*) and the production of the cytolytic toxin Candidalysin (*ECE1*) (**Figure 2C**). Coactivation of morphogenesis genes in *C. albicans* together with iron uptake and utilization was previously observed and might reflect the adaptive prediction phenomenon (Tagkopoulos et al., 2008; Mitchell et al., 2009; Roy and Kornitzer, 2019). This concept suggests that as fungal cells invade host cells, metabolic processes such as metal uptake and utilization are anticipatedly activated to accommodate the metabolic demands of fungal cells. Alternatively, the upregulation of Fe regulon might reflect a situation where this essential metal is depleted as a consequence of Mn starvation. Indeed, ICP-MS quantification showed that the abundance of Fe was significantly reduced in *C. albicans* cells under Mn limitation as compared to Mn-replete conditions (**Figure 2E**). Thus, Mn depletion led to Fe deficiency in *C. albicans* as was observed in bacteria, plants and algae (Tanaka and Navasero, 1966; Allen et al., 2007; Puri et al., 2010; Helmann, 2014). Furthermore, our ICP-MS data also show that Zn levels, but not Cu, were reduced which might also explain the up-regulation of the Zn transporter Zrt101 at 90 min growth in SC-Mn (**Figure 2C and Supplementary Table S1**). These results indicate that *C. albicans* Fe and Zn homeostasis are modulated by intracellular Mn levels.

**Figure 2.**
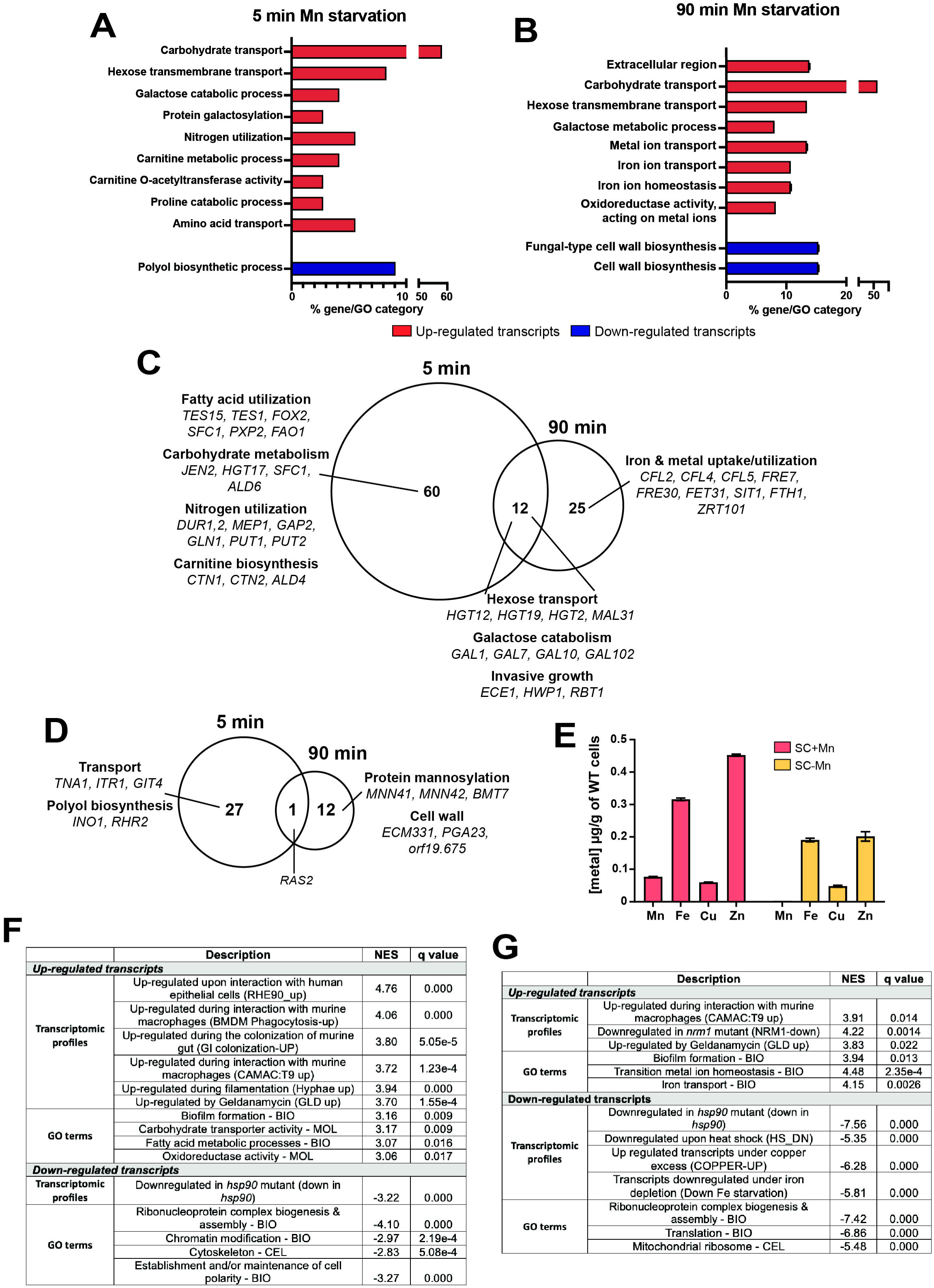
Transcriptomic analysis of *C. albicans* response to Mn starvation. (**A-B**) Gene ontology enrichment of upregulated and downregulated transcripts of *C. albicans* cells growing under Mn limitation compared to cells growing under Mn repletion at 5 min (**A**) and 90 min (**B**). (**C**-**D**) Venn diagram showing overlaps between transcripts upregulated (**C**) or downregulated (**D**) under Mn depletion at 5 min and 90 min. Relevant biological functions and their associated genes are shown. (**E**) Quantification by ICP-MS of Fe, Mn, Zn and Cu in exponentially grown WT cells in SC-Mn and SC+Mn media at 30°C. (**F**-**G**) GSEA analysis of the *C. albicans* Mn-modulated transcriptome at 5 min (**F**) and 90 min (**G**). The vertical black lines indicate the position of each of the genes modulated by Mn in the three ordered data sets (GLD_UP: Transcripts upregulated by geldanamycin; Down in HSP90: Transcripts downregulated in the repressible mutant Tet-HSP90 and HS_DN: transcripts downregulated by heat). The green curve represents the NES (normalized enrichment score) curve, which is the running sum of the weighted enrichment score obtained from GSEA software. NES and nominal *q*-value obtained from the GSEA analysis are shown on the bottom of each plot. The complete GSEA correlations are listed in **Supplementary Table S2**. NES, normalized enrichment score; False Discovery Rate (q-value) of 1%.

To further explore the biological processes modulated by Mn abundance, we used gene set enrichment analysis (GSEA) to elucidate resemblance with the set of the previously published *C. albicans* transcriptional profiling experiments (Sellam et al., 2012). Upregulated transcripts under Mn starvation exhibit significant similarity with transcriptional profiles reflecting different context of host-*C. albicans* interactions including colonization of mammalian gut, infection of human oral epithelial cells and macrophages (**Figure 2F-G and Supplementary Table S2**). This finding suggests that Mn depletion is a situation that *C. albicans* might encounter *in vivo*. Furthermore, transcriptional programs associated with invasive filamentous growth and biofilm formation were also positively correlated with activated transcripts in Mn-starved cells. Intriguingly, GSEA analysis revealed that both 5- and 90-min Mn starvation transcriptomes were correlated with gene signatures of cells where the essential chaperone Hsp90 was inhibited (cells treated with geldanamycin, transcripts downregulated in *hsp90* mutant, and transcripts repressed by heat) (**Figure 2F-G and Supplementary Table S2**). Together, these results imply that Mn limitation generates a transcriptional signature that is reminiscent of heat stress and the expression of fungal virulence.

### Mn starvation induces the unfolded protein response

Transcript levels of three mannosyltransferases (*MNN41*, *MNN42*, *BMT7*) which are enzymes required for the glycosylation of proteins, a process required for protein secretion and cell wall integrity (Martin Loibl and Sabine Strahl, 2013), were reduced under Mn limiting conditions (**Figure 2D**). Given the requirement of Mn for the activity of these enzymes, this might reflect a reduced mannosyltransferases activity in *C. albicans* cells. First, we tested the impact of Mn abundance on the glycosylation levels of *C. albicans* proteins in the WT strain under both Mn limitation and repletion. Surprisingly, under Mn scarcity, we noticed staining of glycosylated proteins with high molecular weight (>250 kDa) as compared to Mn repletion exhibiting staining of proteins with molecular weight higher than 130 kDa (**Figure 3A**). Under Mn limitation, while the WT strain displayed staining of glycosylated proteins with exclusively high molecular weight (>250 kDa), *smf12* mutant exhibited a staining of proteins with lower molecular weight (< 100 kDa) that is indicative of a decrease in the amount and the length of glycosyl residues linked to proteins (**Figure 3A**). This observed weight shift is related to shortage of intracellular Mn as repletion led to a profile indistinguishable from that of the WT strain. Furthermore, our RNA-seq analysis of *C. albicans* cells experiencing Mn starvation revealed a transcriptional signature similar to that of cells where Hsp90 is either genetically or pharmacologically compromised (**Figure 3B**). As Hsp90 primary role is to ensure correct folding and stability of proteins, we hypothesized that upon Mn depletion this functionality is requested as a part of the unfolded protein response (UPR) triggered by a decrease of protein glycosylation as previously reported (Martin Loibl and Sabine Strahl, 2013). In eukaryotic cells, UPR is a signaling pathway activated by multiple endoplasmic reticulum (ER) stresses to preserve protein homeostasis (Hetz, 2012). ER stress is sensed by the Ire1, an ER-located transmembrane endoribonuclease that excises an unconventional intron from the transcription factor *HAC1* mRNA which lead to its activation and induction of the UPR transcriptional response (Wu et al., 2014). To test whether Mn starvation activates UPR response in *C. albicans*, we used RT-PCR to assess the splicing variants of *HAC1* as a proxy of Ire1 activity in cells exposed to Mn starvation at different time-points (0, 5, 30 min). WT cells growing for 30 min under Mn limitation exhibited an increased level of the spliced form of *HAC1* as compared to the repleted condition (**Figure 3C-D**). Splicing of *HAC1* was more marked in *smf12* mutant in absence of Mn and almost entirely turned off at 30 min of growth in the repleted medium. We also found that *ire1* mutant exhibited a glycosylation pattern similar to that of *smf12* when Mn was omitted from the growth medium, however, *ire1* was insensitive to Mn supplementation as the glycosylation pattern remained unchanged (**Figure 3A**). These data suggest that Mn starvation induces UPR response that is signaled by the Ire1-Hac1 regulatory axis. Accordingly, growth alleviation by Mn supplementation observed in WT strain under Mn starvation was not perceived in either *ire1* or *hac1* mutants (**Figure 3E**). Recently, UPR signaling pathway was also shown to be essential for iron uptake in *C. albicans* by controlling localization of the high-affinity iron permease Ftr1 to the cell membrane (Ramírez-Zavala et al., 2022). Thus, UPR signaling appears to play an important role for both Fe and Mn homeostasis in *C. albicans*.

**Figure 3.**
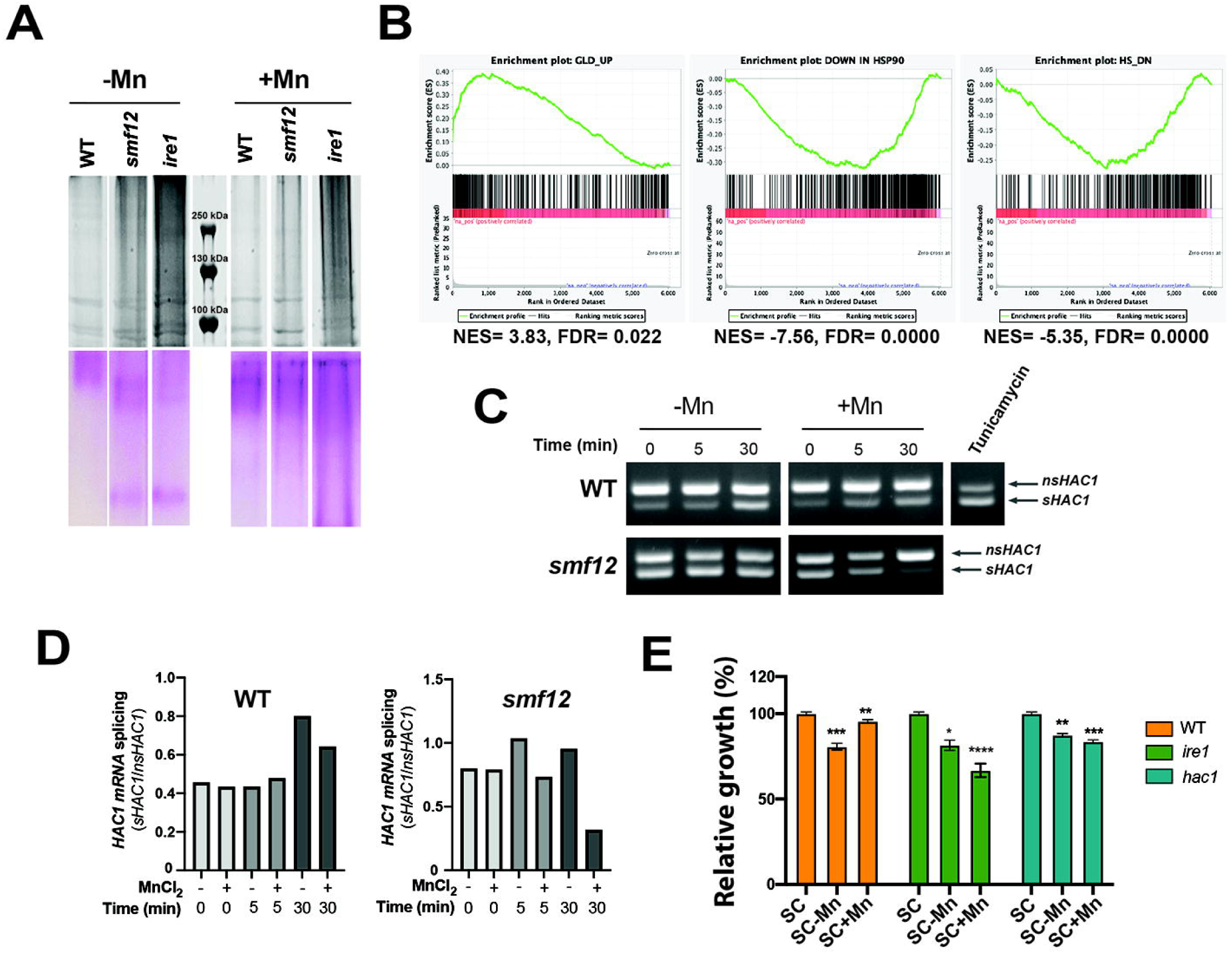
Mn limitation promotes the unfolded protein response in *C. albicans*. (**A**) Impact of *C. albicans* Mn homeostasis on protein glycosylation. *C. albicans* cells were grown as for the RNA-seq experiment and proteins were analyzed on SDS-PAGE with silver stain (upper panel). The same samples on SDS-PAGE were stained using the Glycoprotein Staining Kit (lower panel). (**B**) GSEA graphs of significant correlations between the Mn-depletion transcriptome and Hsp90 inhibition. (**C-D**) Mn starvation induces *HAC1* mRNA splicing. WT and *smf12* cells were grown in SC-Mn at the indicated time and HAC1 splicing was assessed using RT-PCR (**C**). The glycosylation inhibitor and the UPR inducer tunicamycin, was used as a positive control. *nsHAC1*, non-spliced *HAC1*; *sHAC1*, spliced *HAC1*. (**D**) Quantitative assessment of *HAC1* splicing. The intensity of PCR bands was measured using ImageJ and data are presented as the ratio of the sHAC1/nsHAC1 PCR band intensity. (**E**) Impact of Mn depletion on the growth of *ire1* and *hac1* mutants. *C. albicans* strains were grown at 30°C for 13 hrs in SC-Mn or SC+Mn. Growth was normalized to the SC condition. Statistics are multiple t-test with **** p-value < 0.00001, *** p-value < 0.0001, ** p-value < 0.001, * p-value < 0.05.

### Mn modulates host invasion and interaction with immune cells

As the Mn-starvation transcriptome was similar to that expressed during *C. albicans* interaction with different host cells, we wanted to test the impact of Mn homeostasis on the ability of this yeast to damage the host. We found that *C. albicans* cells pre-grown in Mn-starved medium cause more damage to both human enterocytes and murine macrophages than those thriving in Mn repleted medium (**Figure 4A**). The ability of *smf12* mutant to damage HT-29 enterocytes was significantly impaired as compared to the parental WT or the complemented strains regardless of Mn levels in fungal pre-cultures (**Figure 4B**). However, when co-incubated with the J774.A.1 murine macrophage, *smf12* mutant exhibited a similar damage level as the WT strain (**Figure 4C**). We also used the *Galleria mellonella* larvae-*C. albicans* model of systemic candidiasis to assess the impact of Mn levels in fungal pre-cultures on the infectivity of both WT and *smf12* strains. At the first day of infection, *C. albicans* WT pre-grown under Mn sufficiency resulted in the death of 80% of *Galleria* larvae, whereas those cultured in Mn-depleted medium caused 95 % death (**Figure 4D**).While *smf12* mutant pre-grown under Mn replete condition exhibited a similar infectivity rate as the WT strain, *smf12* pre-cultured under Mn limiting condition exhibited an attenuated virulence with a mortality rate of 5% and 50 % at day 1 and 5, respectively, as compared to WT strain that led to 100% mortality at day 1 (**Figure 4D**). Together, these findings suggest that Mn starvation promotes fungal invasiveness as was shown for other essential metals, and that Smf12-mediated Mn homeostasis contributes to fungal virulence.

**Figure 4.**
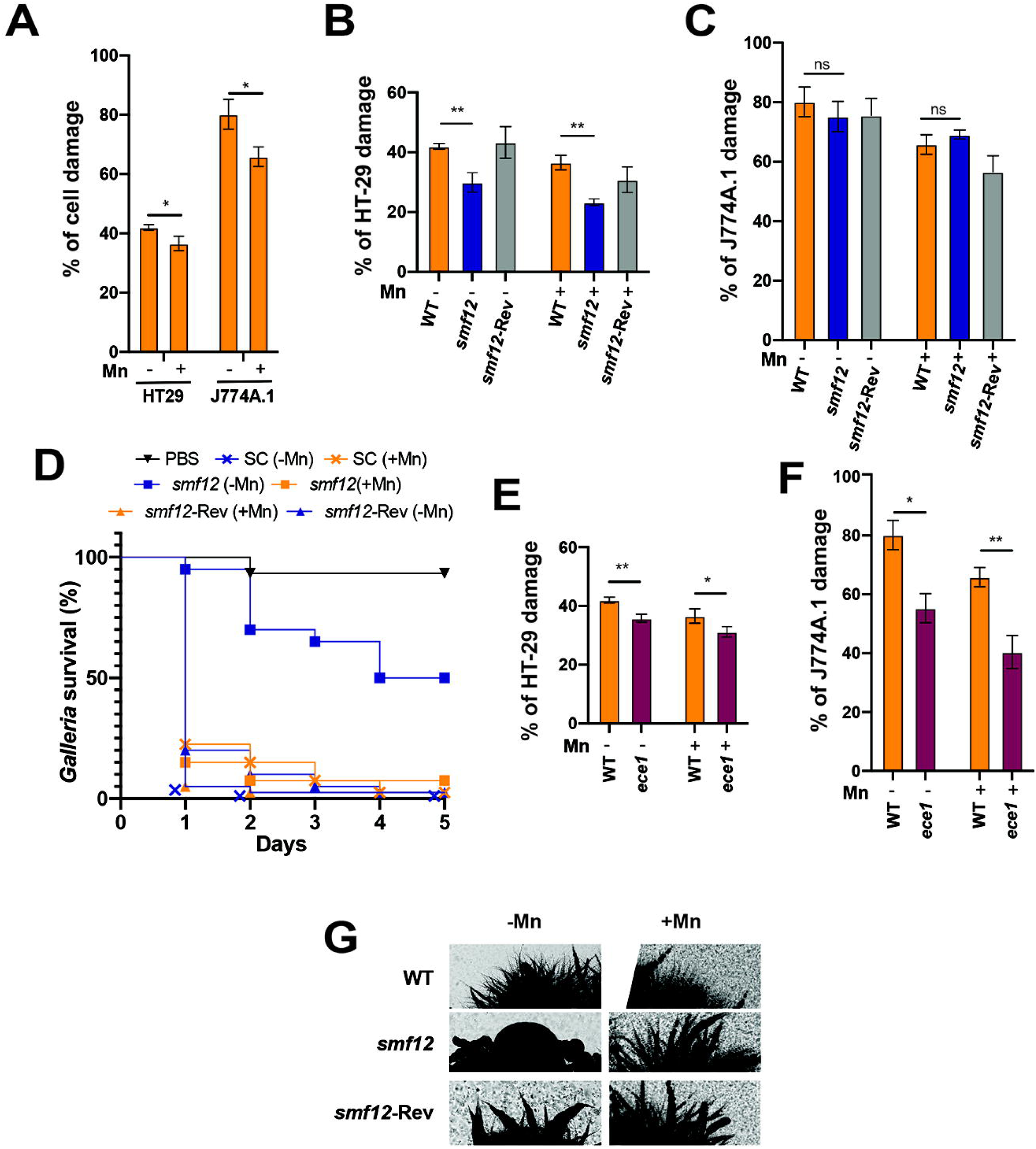
Mn abundance modulate fungal virulence. (**A-C**) The effect of Mn abundance in pre-culture medium on *C. albicans* WT (**A**) and *smf12* infectivity of HT-29 human enterocytes (**B**) and J774A.1 murine macrophages (**C**). Cell damage was assessed using the LDH release assay and was calculated as a percentage of LDH activity as described in the method section. (**D**) Impact of Mn levels in *C. albicans* pre-culture medium and *smf12* mutation on *G. mellonella* larvae infection. WT, *smf12* and *smf12* complemented (*smf12*- Rev) strains, together with the PBS control, were injected into *G. mellonella* larvae and survival was monitored daily for a period of 5 days. (**E**-**F**) Effect of Mn levels on the expression of candidalysin. WT and *ece1* strains pre-cultured in either Mn-depleted or repleted growth media were tested for their ability to cause damage to HT-29 enterocytes and J774A.1 macrophages. Statistics are multiple t-test with ** p-value < 0.001, * p-value < 0.05, ns: non-significant. (**G**) *smf12* morphogenetic defect. Colony morphologies of the WT, *smf12* and the complemented strain (*smf12*-Rev) after 3 days at 37°C on solid Spider medium under both Mn scarcity and repletion.

As the *ECE1* gene encoding candidalysin was upregulated under Mn limitation, we assessed whether the enhanced damage to host cells under this condition was mediated by the activity of this fungal cytolytic toxin. Consistent with previous work, *ece1* mutant caused less damage as compared to the WT strain (Moyes et al., 2016), however, the enhanced damage reported under Mn limitation was also perceived in *ece1* (**Figure 4E-F**). This suggests that Mn modulation of fungal invasiveness is independent of the activity of candidalysin. We did not notice any significant difference regarding the ability of *C. albicans* WT strain to form filaments when using hyphae-promoting growth medium with depleted and repleted Mn levels (**Figure 4G**). Nevertheless, in Mn starved medium, *smf12* mutant formed smooth colonies in contrast to the WT or the complemented strains that differentiated marked invasive filaments (**Figure 4G**). This observation might explain the reduced virulence of *smf12* in the different tested infection models and supports the role of Mn homeostasis in promoting morphogenetic switch in *C. albicans*.

### Mn homeostasis modulates sensitivity to antifungals

The impact of Mn homeostasis on antifungal stress was tested in both WT and *smf12* cells, growing in either Mn limitation or sufficiency. Our data show that sensitivity to fluconazole and miconazole in WT cells was moderately increased in Mn-starved medium as compared to Mn-replete condition (**Figure 5A**). Furthermore, *smf12* was hypersensitive to the two tested azoles as compared to WT specifically in Mn-restricted condition (**Figure 5A**). Assessing the number of CFU confirmed a decrease in cell viability for *smf12* as compared to the WT especially for miconazole treatment and under Mn limitation (**Figure 5B**). Thus, in addition to UPR stress, Mn abundance modulates *C. albicans* tolerance to azole antifungals.

**Figure 5.**
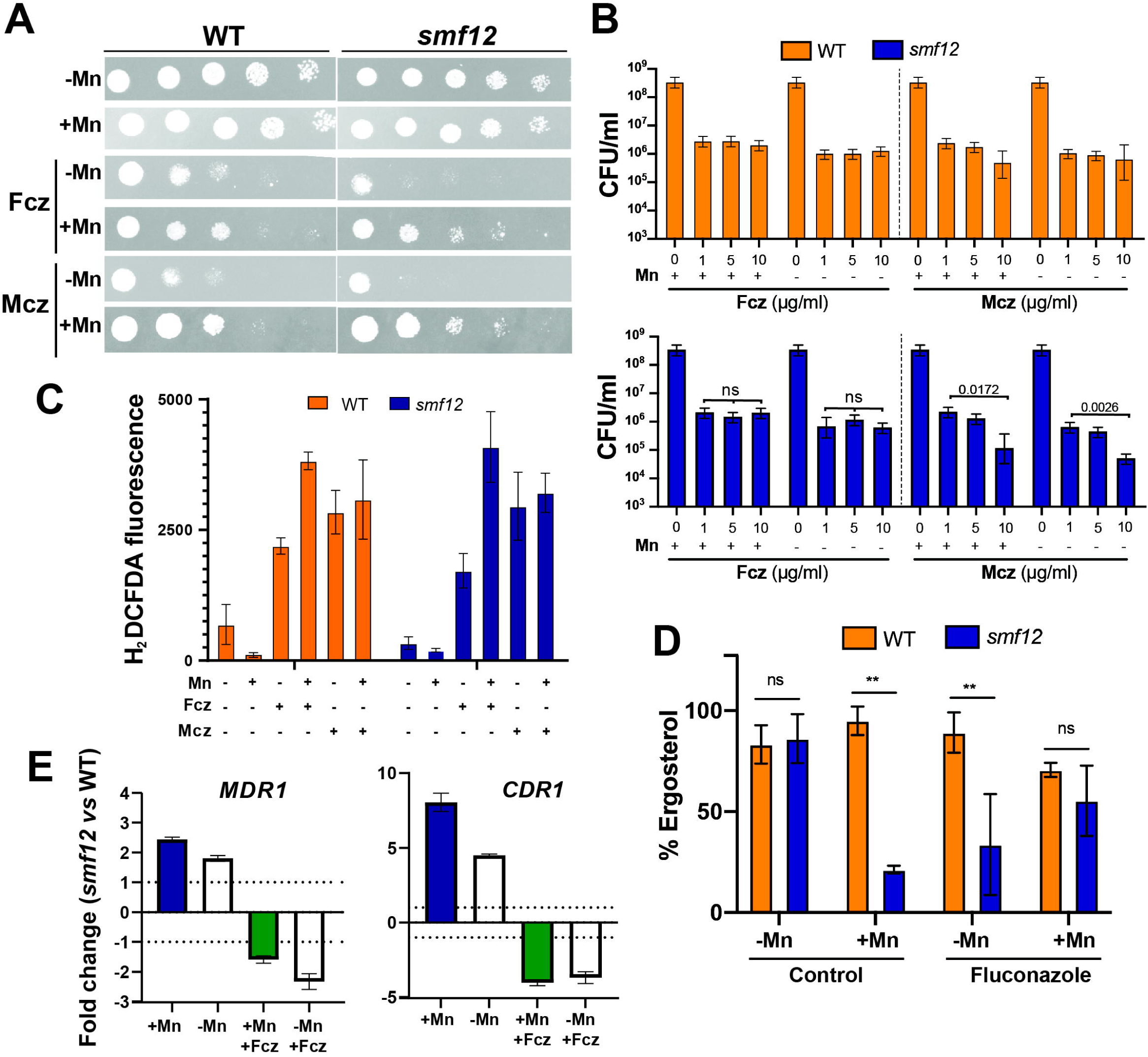
Mn homeostasis modulates antifungal sensitivity. (**A**) Growth assessment of WT and *smf12* mutant cells in the presence of fluconazole (Fcz; 1µg/ml) and miconazole (Mcz; 0.5 µg/ml). *C. albicans* WT and *smf12* strains were serially diluted, spotted on SC-Mn and SC+Mn and incubated for 2 days at 30◦C. (**B**) Evaluation of the impact of Mn availability on WT and *smf12* mutant by CFU counts. Quantification of intracellular ROS (**C**), ergosterol levels (**D**) in WT and *smf12* cells. (**E**) Evaluation of transcript levels of *MDR1* and *CDR1* by qPCR. Transcript levels were assessed under Mn limitation and sufficiency in response to fluconazole and fold-changes were calculated using the comparative ΔCt method. Data were normalized using Ct values obtained from actin in each condition.

To underline the mechanism leading to antifungal sensitivity of either WT or *smf12* mutant under Mn limitation, we first tested whether reactive oxygen species (ROS) as a mediator of azoles antifungal activity was exacerbated (Delattin et al., 2014). As Mn is a critical cofactor of *C. albicans* superoxide dismutases (Rhie et al., 1999; Lamarre et al., 2001), we hypothesized that antifungal sensitivity might be a consequence of impairment of the ROS-neutralizing capacity of *C. albicans* cells. Our data show that neither Mn depletion nor *smf12* mutation has a significant impact on ROS levels in either antifungal-treated or control cells (**Figure 5C**). Secondly, as the cellular content of ergosterol is associated with azole sensitivity (Flowers et al., 2012), we tested the impact of Mn availability and *smf12* mutation on the abundance of this fungal sterol. In WT cells, no perceptible change in ergosterol levels was seen in both fluconazole-treated or non-treated cells under either Mn scarcity or sufficiency (**Figure 5D**). However, for *smf12* mutant, ergosterol amounts dropped significantly in cells growing in SC+Mn medium as compared to SC-Mn. This trend was inverted when *smf12* cells were challenged by fluconazole where ergosterol amount was reduced by approximately 3-fold under Mn starvation as compared to Mn replete condition (**Figure 5D**). Thus, the reduced amount of ergosterol in *smf12* might explain the sensitivity of this mutant to azole antifungals. Lastly, we assessed the contribution of the two drug efflux pumps Cdr1 and Mdr1, which are key determinants of azole clinical resistance (Nishimoto et al., 2019), to the Mn-modulated antifungal sensitivity by assessing their transcript levels using qPCR. Both *MDR1* and *CDR1* were significantly downregulated in *smf12* mutant in *C. albicans* challenged with fluconazole (**Figure 5E**). Taken together, Mn modulation of antifungal sensitivity might operate through the regulation of antifungal efflux and the maintenance of ergosterol homeostasis.

### Global transcriptional response *of C. albicans* to Mn excess

While increasing copper levels is thought to be an *in vivo* defense strategy to limit fungal infections by the host, nothing is known regarding the contribution of Mn to such a mechanism (Mackie et al., 2016). Recent study uncovered a significant increase of Mn in response to systemic infection by *C. albicans* or under colitis (Sunuwar et al., 2023). However, the relevance of such a phenomenon as an antifungal defense mechanism remains unexplored. We undertook RNA-seq profiling of *C. albicans* cells exposed to Mn excess to understand how fungal cells cope with such stress. The inhibitory effect of Mn excess on *C. albicans* growth was detected at 10 mM leading to 10% inhibition while 15 mM Mn caused 70 % growth reduction (**Figure 6A**). To capture the cellular processes impacted by Mn excess, we used RNA-seq profiling of cells exposed to 7.5 mM MnCl_2_, a concentration that led to 10% growth inhibition. Gene ontology enrichment analysis of upregulated transcripts uncover a transcriptional signature of ER stress with enrichment of many biological functions including proteolysis, vesicle-mediated transport, protein folding and response to oxidative stress which are *bona fide* hallmarks of UPR (**Figure 6B and Supplementary Table S1**). Transcripts related to ribosome biogenesis and ribosomal RNA processing were repressed. UPR activation by Mn excess was supported by Hac1 splicing and the requirement of Ire1 to tolerate higher concentrations of Mn (**Figure 6C-E**). This finding implies that both Mn starvation and excess in *C. albicans* generate ER stress which promotes the activation of UPR signaling.

**Figure 6.**
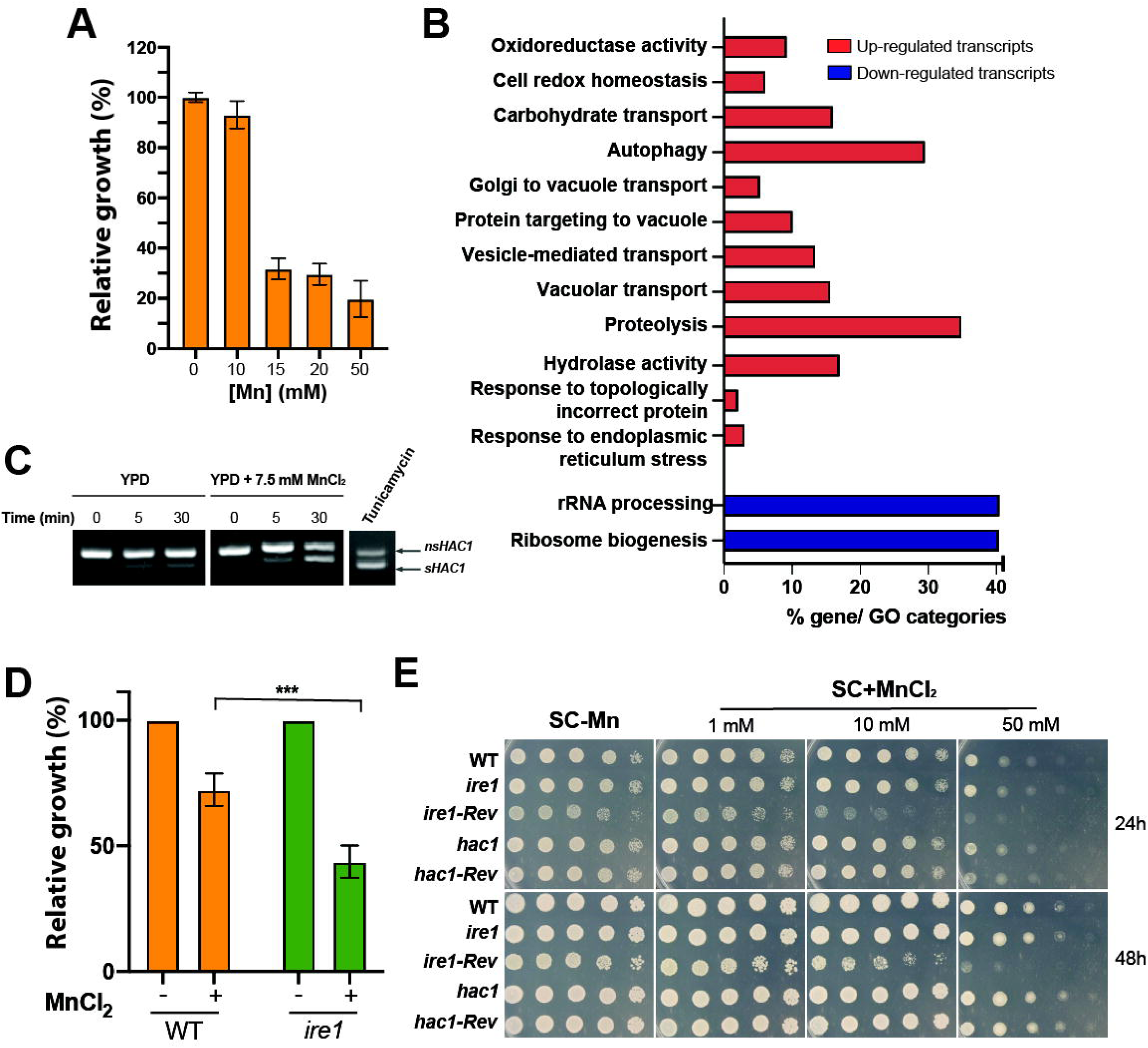
Impact of Mn excess on *C. albicans*. (**A**) Effect of increased Mn levels on *C. albicans* growth. SC5314 was grown in YPD supplemented with different concentrations of MnCl_2_ for 13 h at 30°C. Growth was normalized to the non-treated condition. (**B**) Transcriptional profiling of *C. albicans* cells under Mn excess. Overrepresented GO functional categories of differentially modulated transcripts are shown. (**C**) Mn toxicity induces *HAC1* mRNA splicing. WT cells were grown in YPD supplemented with 7.5 mM of MnCl_2_ and *HAC1* splicing was assessed using RT-PCR. Tunicamycin was used as a positive control. *nsHAC1*, non-spliced *HAC1*; *sHAC1*, spliced *HAC1*. Effect of Mn excess on *ire1* and *hac1* growth in both liquid (**D**) and solid medium (**E**). All strains were grown in liquid YPD with 7.5 mM of MnCl_2_ for 13 hrs at 30°C (**D**) or in YPD-agar medium with the indicated Mn concentrations (**E**). *ire1-Rev* and *hac1-Rev* are the *ire1* and *hac1* complemented strains, respectively. Statistics are multiple t-test with *** p-value < 0.0001.

## Conclusion

Overall, the current study presents a comprehensive transcriptional portrait of biological functions that are modulated by Mn abundance in *C. albicans* cells. Some of these functions have been previously shown to be essential for host infection underscoring the value of targeting fungal Mn homeostasis for potential antifungal therapeutics development. This was also supported by our finding showing that inactivation of the Mn transporter *SMF12* led to attenuated virulence in different infection models. Furthermore, the transcriptional pattern of Mn starvation was similar to that expressed in different contexts of *C. albicans*-host interaction emphasizing that Mn scarcity is a situation encountered during infection. Importantly, we found that intracellular Mn levels influence the abundance of the essential metals Fe and Zn which emphasizes the complex crosstalk between metal ions. This could be explained by the fact that these metals might share common transporters as was reported for NRAMP transporters and their dual specificity toward Mn and Fe (Grotz and Guerinot, 2006; Rai et al., 2021). Alternatively, this might be related to the fact that Fe and Zn assimilation are mediated by proteins that are glycosylated in the secretory pathway, a process that relies on Mn-dependent glycosyltransferase (Zhao et al., 2014; Thines et al., 2018; Thomine and Merlot, 2021). Overall, this work provides fertile areas for future studies to examine the link between Mn metabolism and essential functions that modulate fungal virulence and fitness inside the host.

## Materials and methods

### Strains and growth conditions

The fungal strains used in this study are listed in **Supplementary Table S3**. *C. albicans* clinical strain SC5314 (Fonzi and Irwin, 1993) and its derivatives were routinely maintained at 30°C on synthetic complete (SC; 1.7% yeast nitrogen base, 0.5% ammonium sulfate, 2% dextrose, 0.2% amino acid, with 50 µg/ml uridine) or YPD media (2% Bacto peptone, 1% yeast extract, 2% dextrose, and 50 µg/ml uridine). *smf12* deletion mutant (*smf12*:*URA3*/*smf12*:*HIS1*) was constructed from SN148 strain (Noble and Johnson, 2005) by replacing the entire ORF by a PCR- disruption cassette generated from pFA plasmids (Gola et al., 2003). For complementation of *smf12* mutant, PCR primers were designed to amplify 1kb upstream of *SMF12* in addition to the complete *SMF12* ORF. The resulting PCR products were cloned in the pDUP3 plasmid (Gerami-Nejad et al., 2013). The resulting pDUP3-*SMF12* construct was digested by *SfI1* and integrated into the *NEU5L* genomic site of the *smf12* strain as previously described (Gerami-Nejad et al., 2013) using lithium acetate transformation procedure (Wilson et al., 2000). Transformants were selected on YPD plates supplemented with 200 μg/ml nourseothricin and correct integration was verified by PCR. Primers used for *SMF12* cloning in pDUP3 plasmid and for the diagnosis of integration are listed in **Supplementary Table S3**. Effect of Mn abundance on fungal growth was performed as follows: fungal inocula were prepared from an overnight culture grown in SC medium at 30°C and diluted to a starting OD_600_ of 0.1 in either SC without manganese (SC-Mn) prepared using yeast nitrogen base without manganese (Formidium™) or supplemented with 1 mM of MnCl_2_ (SC+Mn). Cultures were exponentially grown for 13 hours at 30°C under agitation. Relative growth was assessed using the growth on SC medium as a control condition. Effect of Mn excess was tested similarly using YPD medium with different concentration of MnCl_2_ and growth was normalized to the non-treated condition. The filamentation assay was performed in spider medium (Liu et al., 1994). Exponentially growing *C*. *albicans* cells were seeded on Spider- agar plate and incubated for 3 days at 37 °C. All chemicals used in this study were provided by Sigma-Aldrich (St. Louis, MO, United States). Miconazole (1 mg/ml) and fluconazole (10 mg/ml) stock solutions were prepared using dimethyl sulfoxide (DMSO). Working stock solutions of Manganese (II) Chloride (MnCl_2_; 1 M) were prepared using Chelex resin-treated MilliQ water. For growth inhibition assays in liquid SC-Mn and SC+Mn media, overnight cultures of *C. albicans* were resuspended in fresh SC-Mn medium at an OD_600_ of 0.1 and added to a flat-bottom 96-well plate in a total volume of 200 μl per well along with the compounds being tested. For each experiment, a compound-free positive growth control and a cell-free negative control were included. Growth assay curves were performed in triplicate in 96-well plates using a Sunrise plate-reader (Tecan) at 30°C under constant agitation. The CFU assay was performed as following: *C. albicans* cells were grown in presence of 1 ug/ml, 5ug/ml or 10 ug/ml Fluconazole or Miconazole during 48h at 30°C in 96-well plates using the Sunrise plate-reader (Tecan) under constant agitation. Plates were then centrifuged, washed with PBS, and spread on YPD plates at different dilutions. CFU were assessed after 24h growth at 30°C.

### Inductively coupled plasma-mass spectrometry (ICP-MS)

The total amounts of cell-associated Mn were quantified from cultures that had been grown exponentially to an OD_600_ of 0.5 on SC-Mn and SC+Mn. All vessels of yeast culture were washed with 10% nitric acid. Cells were centrifuged, washed with ice-cold metal-free PBS pH 7.4 and dried overnight at 65°C. Dried cells were weighed prior to digestion in a mix of 1 ml NHO_3_ 70% and 2 ml H_2_O_2_ 30% and heated 45 min at 95°C in a dry bath. Digested samples were diluted with 1% NHO_3_. Elemental composition was analyzed by Thermo Scientific iCAP Q ICP-MS instrument (Laser-ablation ICP-MS Facility, Department of Earth and Planetary Sciences, McGill University). Metal concentrations were calculated from the standard curve of Mn (1-800 ppb Mn) and normalized to pellet weight of each sample.

### Expression analysis by RNA-seq and quantitative PCR

Overnight cultures of SC5314 strain were diluted to an OD_600_ of 0.1 in 60 ml of fresh SC-Mn medium and grown at 30°C under agitation (200 rpm) to early logarithmic phase (OD_600_=0.4). Cultures were then either left untreated or supplemented with 1 mM MnCl_2_ and incubated at 30°C for 5 and 90 min. For RNA-seq profiling of Mn excess, overnight cultures of WT cells were diluted to an OD_600_ of 0.1 in fresh 100 ml YPD and incubated with shaking at 30°C to an OD_600_ of 0.8 and split into 50-ml cultures. MnCl_2_ was added to the experimental culture to a final concentration of 7.5 mM, while an equal volume of sterile water was added to the control culture and incubated for 30 min. Cells were harvested by centrifugation and were flash-frozen and stored at -80°C. For each condition, a total of two biological replicates were considered for RNA-seq analysis. Total RNA was extracted using an RNAeasy purification kit (Qiagen) and glass bead lysis in a Biospec Mini 24 bead-beater as previously described (Tebbji et al., 2014). RNA integrity was assessed using Agilent 4200 Tape Station System prior to cDNA library preparation. The NEBNext Ultra II RNA Library Prep Kit for Illumina was used to construct the RNA-seq library following the manufacturer’s instruction. A 2 × 100 paired-end sequencing of cDNAs was performed using an Illumina NovaSeq 6000 sequencing system. The GSEA (Gene Set Enrichment Analysis) pre-ranked tool (http://www.broadinstitute.org/gsea/) was used to determine statistical significance of correlations between the *C. albicans* Mn-sensitive transcriptomes with GO biological process terms and different omics datasets as described in (Sellam et al., 2012; Burgain et al., 2019). Differentially expressed transcripts in **Supplementary Table S1** were identified using Welch’s t-test with a false-discovery rate (FDR) of 5% and 1.5-fold enrichment cut-off. Gene ontology (GO) analysis was performed using GO Term Finder of the Candida Genome Database (Binkley et al., 2014). All RNA-seq data are available at the GEO database (https://www.ncbi.nlm.nih.gov/geo/) with the accession number GSE245114.

For quantitative PCR experiments of *MDR1* and *CDR1* transcripts, a total of three biological and three assay replicates were performed. Cells were grown exponentially to OD_600_=0.5 and treated with fluconazole (1µg/ml) in either SC-Mn or SC+Mn for 90 min. RNAs were extracted as for the RNA-seq experiment. cDNA was synthesized from 1 μg of total RNA using the High-Capacity cDNA Reverse Transcription kit (Applied Biosystems). The mixture was incubated at 25°C for 10 min, 37°C for 120 min and 85°C for 5 min. RNAse H (NEB) was added to remove RNA and samples were incubated at 37°C for 20 min. qPCR was performed using a StepOnePlus Instrument (Applied Biosystems) for 40 amplification cycles with the PowerUp SYBR Green master mix (Applied Biosystems). The reactions were incubated at 95°C for 10 min and cycled for 40 times at 95°C, 15 s; 60°C, 1 min. Fold-enrichment of each tested transcript was estimated using the comparative ΔΔCt method. To evaluate the gene expression level, the results were normalized using Ct values obtained from Actin (*ACT1*, C1_13700W_A). Primer sequences used for this analysis are summarized in **Supplementary Table S3**.

### HAC1 mRNA splicing assay

The *HAC1* splicing was assessed using reverse transcription-PCR (RT-PCR) as previously described (Chaillot et al., 2015). RNAs were extracted from *C. albicans* cells challenged with tunicamycin (10 μg/ml), used as a positive control, or in the absence or the presence of manganese (1mM MnCl_2_ for Mn repletion and 7.5mM for Mn excess experiments) as described in the RNA-seq experiment. cDNAs were obtained using Superscript II reverse transcriptase (Applied BioSystems) as recommended by the supplier. The obtained cDNA was used as a template to amplify spliced and un-spliced *HAC1* cDNAs using the primer pair described in (Chaillot et al., 2015). The PCR products were resolved on 4% agarose gel. PCR band intensities were quantified with ImageJ (Irving et al., 2007).

### *Galleria* virulence assay

Larvae of *G. mellonella* (Elevages Lisard, Canada) in the instar larval stage of development were used. Overnight cultures of *C. albicans* strains were washed twice and diluted in 20 µl PBS to obtain a quantity of 5x10^5^ cells in 20μl for injection. *G. mellonella* larvae weighing 180 ± 10 mg were injected between the third pair of prothoracic legs. Infected larvae were incubated at 37 °C. Two replicates, each consisting of 20 larvae, were carried out with survival rates measured daily for a period of 5 days. Death was determined based on the lack of response to touch and the inability to right themselves. Kaplan-Meier survival curves were created and compared with the log-rank test (GraphPad Prism 5).

### HT-29 and J774A.1 damage assay

Damage to the human colon epithelial cell line HT-29 (ATCC; HTB-38) and the murine J774A.1 (ATCC TIB-67) macrophages was assessed using a lactate dehydrogenase (LDH) cytotoxicity detection kit*PLUS* (Roche), which measures the release of the LDH enzyme in the growth medium. HT-29 and J774A.1 cells were grown in 96-well plates as monolayers in McCoy’s medium and Dulbecco’s Modified Eagle’s Medium (DMEM), respectively, supplemented with 10% FBS at 2 × 10^4^ cells per well and incubated at 37°C with 5% CO_2_ overnight. Cells were then infected with *C. albicans* cells pre-cultured in the presence (SC+Mn) or the absence of Mn (SC-Mn), at MOI cell:yeast of 1:2 for 24 h at 37°C with 5% CO_2_. Following incubation, 100 μl of supernatant was removed from each experimental well and LDH activity in this supernatant was determined by measuring the absorbance at 490 nm following the manufacturer’s instructions. Cytotoxicity was calculated as follows: % cytotoxicity = (experimental value - low control [untreated cells]) / (high control [lysis buffer] - low control) × 100.

### Glycosylation assay

*C. albicans* cells were grown in SC-Mn and SC+Mn as for the RNA-seq experiments. Cells were harvested by centrifugation and lysed by bead beating in IP150 buffer (50 mM Tris-HCl at pH 7.4, 150 mM NaCl, 2 mM MgCl_2_, 1% Nonidet P-40, 5% glycerol) supplemented with Complete Mini protease inhibitor mixture tablet (Roche Applied Science) and 10 mM phenylmethylsulfonyl fluoride (PMSF). The lysates were then cleared by centrifugation, and protein concentration was estimated using the Bradford assay. A total of 80 μg of proteins were heated for 5 min at 95°C in the 2X Laemmli buffer and loaded in an Acrylamide gel. Gels have been stained with silver to observe the total protein profile or with Pierce™ Glycoprotein Staining Kit (Thermo Fisher) to detect glycoprotein sugar moieties according to the manufacturer’s instruction.

### ROS quantification

Intracellular ROS was measured using the oxidative stress indicator H_2_DCFA-DA dye (50 μg/ml final concentration; Invitrogen, ThermoFisher Scientific) to quantify the level of reactive oxygen species. Cells were grown as described in the qPCR experiment.

### Ergosterol quantification

Ergosterol quantification was performed as previously described (Arthington-Skaggs et al., 1999). Cells were grown exponentially to OD_600_=0.5 and treated with fluconazole (1µg/ml) either SC-Mn or SC+Mn for 16 hours. Cells were then washed, weighted, resuspended in ethanolic potassium hydroxide at 25%, and incubated for 2 hours in a 95°C water bath. After cooling, 3 ml n-heptane (Heptane, 99%) and 1 ml H_2_O were added. Fluorescence of the supernatants was measured at 282 and 230 nm.

## Authors’ contributions

Conceptualization: MH and AS; Methodology: MH, IK, AV, FT, RA; Funding acquisition: AS; Resources: AS; Supervision: AS; Writing original draft: AS and MH; Writing, review & editing: AS, MH, IK, FT, AV, and RA. All authors listed gave final approval for publication.

## Funding

This work was supported by funds from the Canadian Institutes for Health Research (CIHR project grant), the Canadian Foundation for Innovation and the start-up funds from the Montreal Heart Institute (MHI) to AS. AS is a recipient of the Fonds de Recherche du Québec-Santé (FRQS) Senior salary award. Manon Henry is supported by a PhD fellowship from the Natural Sciences and Engineering Research Council of Canada-CREATE program EvoFunPath. RA is a recipient of the Fonds de recherche du Québec (FRQ) and the Palestine Academy for Science and Technology (PALAST) collaborative fund.

## Availability of data and materials

The original contributions presented in the study are included in the Supplementary Material. RNA-seq data have been submitted to the GEO database at https://www.ncbi.nlm.nih.gov/geo/query/acc.cgi?acc=GSE245114 under accession number GSE245114. Further inquiries can be directed to the corresponding author.

## Conflict of Interest

The authors declare that the research was conducted in the absence of any commercial or financial relationships that could be construed as a potential conflict of interest.

## Acknowledgment

We are grateful to Bernardo Ramírez-Zavala and Joachim Morschhäuser (University of Würzburg) for providing *ire1* and *hac1* mutant strains.

## Notes

### Competing Interest Statement

The authors have declared no competing interest.

## References

Allen, M. D., Kropat, J., Tottey, S., Del Campo, J. A., and Merchant, S. S. (2007). Manganese deficiency in Chlamydomonas results in loss of photosystem II and MnSOD function, sensitivity to peroxides, and secondary phosphorus and iron deficiency. Plant Physiol 143, 263–277. doi: 10.1104/pp.106.088609.

Arthington-Skaggs, B. A., Jradi, H., Desai, T., and Morrison, C. J. (1999). Quantitation of Ergosterol Content: Novel Method for Determination of Fluconazole Susceptibility of Candida albicans. J Clin Microbiol 37, 3332–3337. Available at: https://www.ncbi.nlm.nih.gov/pmc/articles/PMC85559/ [Accessed August 22, 2023].

Bairwa, G., Hee Jung, W., and Kronstad, J. W. (2017). Iron acquisition in fungal pathogens of humans. Metallomics 9, 215–227. doi: 10.1039/C6MT00301J.

Besold, A. N., Gilston, B. A., Radin, J. N., Ramsoomair, C., Culbertson, E. M., Li, C. X., et al. (2018). Role of Calprotectin in Withholding Zinc and Copper from Candida albicans. Infect Immun 86, e00779–17. doi: 10.1128/IAI.00779-17.

Binkley, J., Arnaud, M. B., Inglis, D. O., Skrzypek, M. S., Shah, P., Wymore, F., et al. (2014). The Candida Genome Database: the new homology information page highlights protein similarity and phylogeny. Nucleic Acids Res 42, D711–6. doi: 10.1093/nar/gkt1046.

Brophy, M. B., and Nolan, E. M. (2015). Manganese and Microbial Pathogenesis: Sequestration by the Mammalian Immune System and Utilization by Microorganisms. ACS Chem. Biol. 10, 641–651. doi: 10.1021/cb500792b.

Burgain, A., Pic, É., Markey, L., Tebbji, F., Kumamoto, C. A., and Sellam, A. (2019). A novel genetic circuitry governing hypoxic metabolic flexibility, commensalism and virulence in the fungal pathogen Candida albicans. PLOS Pathogens 15, e1007823. doi: 10.1371/journal.ppat.1007823.

Cai, Z., Du, W., Zeng, Q., Long, N., Dai, C., and Lu, L. (2017). Cu-sensing transcription factor Mac1 coordinates with the Ctr transporter family to regulate Cu acquisition and virulence in Aspergillus fumigatus. Fungal Genet. Biol. 107, 31–43. doi: 10.1016/j.fgb.2017.08.003.

Chaillot, J., Tebbji, F., Remmal, A., Boone, C., Brown, G. W., Bellaoui, M., et al. (2015). The Monoterpene Carvacrol Generates Endoplasmic Reticulum Stress in the Pathogenic Fungus Candida albicans. Antimicrob Agents Chemother 59, 4584–4592. doi: 10.1128/AAC.00551-15.

Citiulo, F., Jacobsen, I. D., Miramón, P., Schild, L., Brunke, S., Zipfel, P., et al. (2012). Candida albicans Scavenges Host Zinc via Pra1 during Endothelial Invasion. PLOS Pathogens 8, e1002777. doi: 10.1371/journal.ppat.1002777.

Corbin, B. D., Seeley, E. H., Raab, A., Feldmann, J., Miller, M. R., Torres, V. J., et al. (2008). Metal chelation and inhibition of bacterial growth in tissue abscesses. Science 319, 962– 965. doi: 10.1126/science.1152449.

Culbertson, E. M., Bruno, V. M., Cormack, B. P., and Culotta, V. C. (2020). Expanded role of the Cu-sensing transcription factor Mac1p in Candida albicans. Mol Microbiol 114, 1006– 1018. doi: 10.1111/mmi.14591.

Culotta, V. C., Yang, M., and Hall, M. D. (2005). Manganese Transport and Trafficking: Lessons Learned from Saccharomyces cerevisiae. Eukaryotic Cell 4, 1159–1165. doi: 10.1128/ec.4.7.1159-1165.2005.

Cyert, M. S., and Philpott, C. C. (2013). Regulation of Cation Balance in *Saccharomyces cerevisiae*. Genetics 193, 677–713. doi: 10.1534/genetics.112.147207.

Delattin, N., Cammue, B. P., and Thevissen, K. (2014). Reactive oxygen species-inducing antifungal agents and their activity against fungal biofilms. Future Medicinal Chemistry 6, 77–90. doi: 10.4155/fmc.13.189.

Flowers, S. A., Barker, K. S., Berkow, E. L., Toner, G., Chadwick, S. G., Gygax, S. E., et al. (2012). Gain-of-function mutations in UPC2 are a frequent cause of ERG11 upregulation in azole-resistant clinical isolates of Candida albicans. Eukaryot Cell 11, 1289–1299. doi: 10.1128/EC.00215-12.

Fonzi, W. A., and Irwin, M. Y. (1993). Isogenic strain construction and gene mapping in Candida albicans. Genetics 134, 717–728. Available at: http://www.ncbi.nlm.nih.gov/pubmed/8349105.

Fourie, R., Kuloyo, O. O., Mochochoko, B. M., Albertyn, J., and Pohl, C. H. (2018). Iron at the Centre of Candida albicans Interactions. Frontiers in Cellular and Infection Microbiology 8. Available at: https://www.frontiersin.org/articles/10.3389/fcimb.2018.00185 [Accessed August 18, 2023].

Gerami-Nejad, M., Zacchi, L. F., McClellan, M., Matter, K., and Berman, J. (2013). Shuttle vectors for facile gap repair cloning and integration into a neutral locus in Candida albicans. *Microbiology (Reading*, Engl.) 159, 565–579. doi: 10.1099/mic.0.064097-0.

Gerwien, F., Skrahina, V., Kasper, L., Hube, B., and Brunke, S. (2018). Metals in fungal virulence. FEMS Microbiology Reviews 42. doi: 10.1093/femsre/fux050.

Gola, S., Martin, R., Walther, A., Dunkler, A., and Wendland, J. (2003). New modules for PCR- based gene targeting in Candida albicans: rapid and efficient gene targeting using 100 bp of flanking homology region. Yeast 20, 1339–1347. doi: 10.1002/yea.1044.

Grotz, N., and Guerinot, M. L. (2006). Molecular aspects of Cu, Fe and Zn homeostasis in plants. Biochimica et Biophysica Acta (BBA) - Molecular Cell Research 1763, 595–608. doi: 10.1016/j.bbamcr.2006.05.014.

Haas, H. (2012). Iron – A Key Nexus in the Virulence of Aspergillus fumigatus. Frontiers in Microbiology 3. Available at: https://www.frontiersin.org/articles/10.3389/fmicb.2012.00028 [Accessed August 18, 2023].

Helmann, J. D. (2014). Specificity of Metal Sensing: Iron and Manganese Homeostasis in Bacillus subtilis. J Biol Chem 289, 28112–28120. doi: 10.1074/jbc.R114.587071.

Hennigar, S. R., and McClung, J. P. (2016). Nutritional Immunity: Starving Pathogens of Trace Minerals. American Journal of Lifestyle Medicine 10, 170–173. doi: 10.1177/1559827616629117.

Hetz, C. (2012). The unfolded protein response: controlling cell fate decisions under ER stress and beyond. Nat Rev Mol Cell Biol 13, 89–102. doi: 10.1038/nrm3270.

Hood, M. I., and Skaar, E. P. (2012). Nutritional immunity: transition metals at the pathogen-host interface. Nat Rev Microbiol 10, 525–37. doi: 10.1038/nrmicro2836.

Hwang, C.-S., Baek, Y.-U., Yim, H.-S., and Kang, S.-O. (2003). Protective roles of mitochondrial manganese-containing superoxide dismutase against various stresses in Candida albicans. Yeast 20, 929–941. doi: 10.1002/yea.1004.

Irving, B. A., Weltman, J. Y., Brock, D. W., Davis, C. K., Gaesser, G. A., and Weltman, A. (2007). NIH ImageJ and Slice-O-Matic Computed Tomography Imaging Software to Quantify Soft Tissue. Obesity 15, 370–376. doi: 10.1038/oby.2007.573.

Jensen, L. T., Ajua-Alemanji, M., and Culotta, V. C. (2003). The Saccharomyces cerevisiae High Affinity Phosphate Transporter Encoded by PHO84 Also Functions in Manganese Homeostasis. Journal of Biological Chemistry 278, 42036–42040. doi: 10.1074/jbc.M307413200.

Khemiri, I., Tebbji, F., and Sellam, A. (2020). Transcriptome analysis uncovers a link between copper metabolism, and both fungal fitness and antifungal sensitivity in the opportunistic yeast Candida albicans. Front. Microbiol. 11. doi: 10.3389/fmicb.2020.00935.

Lamarre, C., LeMay, J.-D., Deslauriers, N., and Bourbonnais, Y. (2001). Candida albicans Expresses an Unusual Cytoplasmic Manganese-containing Superoxide Dismutase (SOD3 Gene Product) upon the Entry and during the Stationary Phase. Journal of Biological Chemistry 276, 43784–43791. doi: 10.1074/jbc.M108095200.

Liu, H., Köhler, J., and Fink, G. R. (1994). Suppression of hyphal formation in Candida albicans by mutation of a STE12 homolog. Science 266. doi: 10.1126/science.7992058.

Liu, N.-N., Uppuluri, P., Broggi, A., Besold, A., Ryman, K., Kambara, H., et al. (2018). Intersection of phosphate transport, oxidative stress and TOR signalling in Candida albicans virulence. PLOS Pathogens 14, e1007076. doi: 10.1371/journal.ppat.1007076.

Mackie, J., Szabo, E. K., Urgast, D. S., Ballou, E. R., Childers, D. S., MacCallum, D. M., et al. (2016). Host-Imposed Copper Poisoning Impacts Fungal Micronutrient Acquisition during Systemic Candida albicans Infections. PLoS ONE 11, e0158683. doi: 10.1371/journal.pone.0158683.

Martin Loibl and Sabine Strahl (2013). Protein O-mannosylation: What we have learned from baker’s yeast. Biochimica et Biophysica Acta (BBA) - Molecular Cell Research 1833, 2438–2446. doi: 10.1016/j.bbamcr.2013.02.008.

Martinez-Finley, E. J., Chakraborty, S., and Aschner, M. (2013). “Manganese in Biological Systems,” in Encyclopedia of Metalloproteins, eds. R. H. Kretsinger, V. N. Uversky, and E. A. Permyakov (New York, NY: Springer), 1297–1303. doi: 10.1007/978-1-4614-1533-6_284.

Mitchell, A., Romano, G. H., Groisman, B., Yona, A., Dekel, E., Kupiec, M., et al. (2009). Adaptive prediction of environmental changes by microorganisms. Nature 460, 220–224. doi: 10.1038/nature08112.

Moyes, D. L., Wilson, D., Richardson, J. P., Mogavero, S., Tang, S. X., Wernecke, J., et al. (2016). Candidalysin is a fungal peptide toxin critical for mucosal infection. Nature 532. doi: 10.1038/nature17625.

Murdoch, C. C., and Skaar, E. P. (2022). Nutritional immunity: the battle for nutrient metals at the host–pathogen interface. Nat Rev Microbiol 20, 657–670. doi: 10.1038/s41579-022-00745-6.

Nishimoto, A. T., Sharma, C., and Rogers, P. D. (2019). Molecular and genetic basis of azole antifungal resistance in the opportunistic pathogenic fungus Candida albicans. J Antimicrob Chemother 75, 257–270. doi: 10.1093/jac/dkz400.

Noble, S. M., and Johnson, A. D. (2005). Strains and strategies for large-scale gene deletion studies of the diploid human fungal pathogen Candida albicans. Eukaryot Cell 4, 298–309. doi: 10.1128/EC.4.2.298-309.2005.

Puri, S., Hohle, T. H., and O’Brian, M. R. (2010). Control of bacterial iron homeostasis by manganese. Proceedings of the National Academy of Sciences 107, 10691–10695. doi: 10.1073/pnas.1002342107.

Rai, S., Singh, P. K., Mankotia, S., Swain, J., and Satbhai, S. B. (2021). Iron homeostasis in plants and its crosstalk with copper, zinc, and manganese. Plant Stress 1, 100008. doi: 10.1016/j.stress.2021.100008.

Ramírez-Zavala, B., Krüger, I., Dunker, C., Jacobsen, I. D., and Morschhäuser, J. (2022). The protein kinase Ire1 has a Hac1-independent essential role in iron uptake and virulence of Candida albicans. PLoS Pathog 18, e1010283. doi: 10.1371/journal.ppat.1010283.

Reddi, A. R., Jensen, L. T., and Culotta, V. C. (2009). Manganese Homeostasis in Saccharomyces cerevisiae. Chem Rev 109, 4722–4732. doi: 10.1021/cr900031u.

Rhie, G., Hwang, C.-S., Brady, M., Kim, S.-T., Kim, Y.-R., Huh, W.-K., et al. (1999). Manganese- containing superoxide dismutase and its gene from Candida albicans. Biochimica et Biophysica Acta (BBA) - General Subjects 1426, 409–419. doi: 10.1016/S0304-4165(98)00161-5.

Roy, U., and Kornitzer, D. (2019). Heme-iron acquisition in fungi. Current Opinion in Microbiology 52, 77–83. doi: 10.1016/j.mib.2019.05.006.

Roy, U., Yaish, S., Weissman, Z., Pinsky, M., Dey, S., Horev, G., et al. (2022). Ferric reductase- related proteins mediate fungal heme acquisition. eLife 11, e80604. doi: 10.7554/eLife.80604.

Sellam, A., Tebbji, F., Whiteway, M., and Nantel, A. (2012). A novel role for the transcription factor Cwt1p as a negative regulator of nitrosative stress in Candida albicans. PloS one 7, e43956. doi: 10.1371/journal.pone.0043956.

Sia, A. K., Allred, B. E., and Raymond, K. N. (2013). Siderocalins: Siderophore binding proteins evolved for primary pathogen host defense. Curr Opin Chem Biol 17, 150–157. doi: 10.1016/j.cbpa.2012.11.014.

Soares, L. W., Bailão, A. M., Soares, C. M. de A., and Bailão, M. G. S. (2020). Zinc at the Host– Fungus Interface: How to Uptake the Metal? Journal of Fungi 6, 305. doi: 10.3390/jof6040305.

Sunuwar, L., Tomar, V., Wildeman, A., Culotta, V., and Melia, J. (2023). Hepatobiliary manganese homeostasis is dynamic in the setting of inflammation or infection in mice. The FASEB Journal 37, e23123. doi: 10.1096/fj.202300539R.

Supek, F., Supekova, L., Nelson, H., and Nelson, N. (1996). A yeast manganese transporter related to the macrophage protein involved in conferring resistance to mycobacteria. Proceedings of the National Academy of Sciences 93, 5105–5110. doi: 10.1073/pnas.93.10.5105.

Tagkopoulos, I., Liu, Y.-C., and Tavazoie, S. (2008). Predictive Behavior Within Microbial Genetic Networks. Science 320, 1313–1317. doi: 10.1126/science.1154456.

Tanaka, A., and Navasero, S. A. (1966). Interaction between iron and manganese in the rice plant. Soil Science and Plant Nutrition 12, 29–33. doi: 10.1080/00380768.1966.10431958.

Tebbji, F., Chen, Y., Albert, J. R., Gunsalus, K. T. W., Kumamoto, C. A., Nantel, A., et al. (2014). A Functional Portrait of Med7 and the Mediator Complex in Candida albicans. PLOS Genetics 10, e1004770. doi: 10.1371/journal.pgen.1004770.

Thines, L., Deschamps, A., Sengottaiyan, P., Savel, O., Stribny, J., and Morsomme, P. (2018). The yeast protein Gdt1p transports Mn2+ ions and thereby regulates manganese homeostasis in the Golgi. J Biol Chem 293, 8048–8055. doi: 10.1074/jbc.RA118.002324.

Thomine, S., and Merlot, S. (2021). Manganese matters: feeding manganese into the secretory system for cell wall synthesis. New Phytologist 231, 2107–2109. doi: 10.1111/nph.17545.

Waldron, K. J., Rutherford, J. C., Ford, D., and Robinson, N. J. (2009). Metalloproteins and metal sensing. Nature 460, 823–830. doi: 10.1038/nature08300.

Weatherburn, J. D. C., Deborah A. Traynor, David C. (2000). “Enzymes and Proteins Containing Manganese: An Overview,” in Metal Ions in Biological Systems (CRC Press).

Wildeman, A. S., Patel, N. K., Cormack, B. P., and Culotta, V. C. (2023). The role of manganese in morphogenesis and pathogenesis of the opportunistic fungal pathogen Candida albicans. PLOS Pathogens 19, e1011478. doi: 10.1371/journal.ppat.1011478.

Wilson, R. B., Davis, D., Enloe, B. M., and Mitchell, A. P. (2000). A recyclable Candida albicans URA3 cassette for PCR product-directed gene disruptions. Yeast 16, 65–70. doi: 10.1002/(SICI)1097-0061(20000115)16:1<65::AID-YEA508>3.0.CO;2-M.

Wu, H., Ng, B. S. H., and Thibault, G. (2014). Endoplasmic reticulum stress response in yeast and humans. Biosci Rep 34, e00118. doi: 10.1042/BSR20140058.

Zhao, N., Zhang, A.-S., Worthen, C., Knutson, M. D., and Enns, C. A. (2014). An iron-regulated and glycosylation-dependent proteasomal degradation pathway for the plasma membrane metal transporter ZIP14. Proceedings of the National Academy of Sciences 111, 9175– 9180. doi: 10.1073/pnas.1405355111.

Zygiel, E. M., and Nolan, E. M. (2018). Transition Metal Sequestration by the Host-Defense Protein Calprotectin. Annual Review of Biochemistry 87, 621–643. doi: 10.1146/annurev-biochem-062917-012312.

